# Molecular architecture of glideosome and nuclear F-actin in *Plasmodium falciparum*

**DOI:** 10.1101/2024.04.22.590301

**Authors:** Vojtech Pražák, Daven Vasishtan, Kay Grünewald, Ross G. Douglas, Josie L. Ferreira

## Abstract

Actin-based motility is required for the transmission of malaria sporozoites. While this has been shown biochemically, filamentous actin has remained elusive and has to date never been directly visualised inside the parasite. Using focused ion beam milling and electron cryo-tomography, we studied dynamic actin filaments in unperturbed *Plasmodium falciparum* cells for the first time. This allowed us to dissect the assembly, path and fate of actin filaments during parasite gliding and determine a complete 3D model of F-actin within sporozoites. We show that within the cell, actin assembles into micrometre long filaments, much longer than observed in *in vitro* studies. After their assembly at the parasite’s apical end, actin filaments continue to grow as they are transported down the cell as part of the glideosome machinery, and are disassembled at the basal end in a rate-limiting step. Large pores in the IMC, constrained to the basal end, may facilitate actin exchange between the pellicular space and the cytosol for its recycling and maintenance of directional actin flow for efficient gliding. The data also reveal striking and extensive actin bundles in the nucleus. Implications of these structures for motility and transmission are discussed.

---

The *Plasmodium falciparum* parasite causes the most severe form of malaria in humans^1^. Infection occurs during a bite from an infected mosquito, where sporozoites leave the mosquito salivary glands and are deposited into the skin^2^. Within the skin, sporozoites move rapidly (1-2 μm.s^-1^) and persistently (> 1 hour) to encounter and traverse peripheral blood capillaries. This parasite stage utilises an uncommon form of motility, termed gliding motility. It relies on a specialised, unconventional actomyosin motor system, situated below the plasma membrane, where the myosin powerstroke results in the rearward translocation of actin filaments (F-actin) and associated adhesins^3^. *Plasmodium* requires two highly sequence divergent actin isotypes for its cellular functions, with actin-1 being expressed throughout the life cycle and directly involved in gliding motility. Biochemically, actin-1 monomers assemble into F-actin at rates similar to vertebrate actin isotypes. However, *Plasmodium* F-actin appears to be dynamically unstable *in vitro* and through shrinkage and fragmentation, ultimately result in only very short filaments of approximately 100 nm length^4–7^. Within the parasite, actin filaments have historically been difficult to visualise and the failure of traditional actin labelling tools on this divergent actin, has limited our understanding of dynamics within the cellular context. Recent work in *Plasmodium* and its related apicomplexan *Toxoplasma*, using the filament recognising actin chromobody, revealed localisations of actin filament pools primarily at the front (apical), rear (basal) and nuclear region of motile cells^8–10^. However, resolving these enigmatic actin filaments has proven difficult and an *in vivo* understanding of the arrangement, lengths, journey and fate of filaments in highly motile *Plasmodium* sporozoites remains unclear.

We used Focussed Ion Beam milling (FIB-milling) and electron cryo tomography (cryo-ET) to image actin filaments and other subcellular structures in *Plasmodium falciparum* sporozoites (Fig. S1). Subvolume averaging (SVA) was used to determine the structure and a complete 3D model of F-actin within sporozoites (Fig. 1). Factin was present in all major subcellular compartments (confirmed by SVA of individual filaments/compartments): the pellicular space (the intermembrane space between the plasma membrane and the inner membrane complex and the primary site for gliding machinery), the cytosol, and most remarkably in the nucleus. Surprisingly, and unlike some previous *in vitro* reports, we consistently observed actin filaments longer than 100 nm, some up to 850 nm long (Fig. 1g, S2b). The mean length of 200 ± 140 nm (standard deviation) is likely an underestimate due to some filaments being truncated by FIB-milling. The global F-actin concentration was measured to be 40 ± 7 μM.

**Fig. 1:**
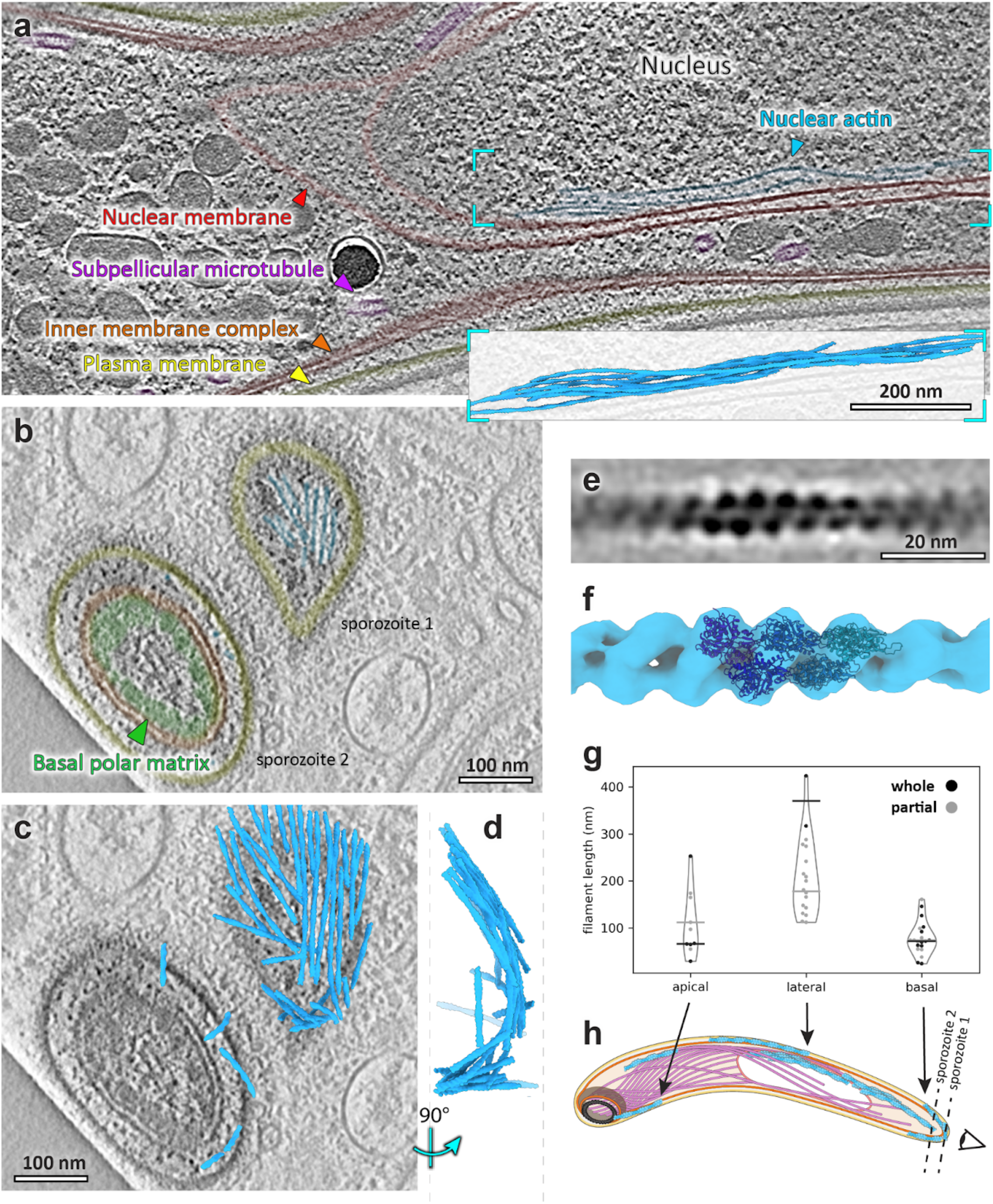
Discrete F-actin populations are found in the sporozoite. **a**, Slice through a tomogram showing a bundle of actin filaments in the nucleus. Insert shows a 3D representation of the actin bundle derived from subvolume averaging. **b**, Slice through the basal ends of two sporozoites (see **h** for positioning of the slicing planes relative to the cell). **c**, Same as **b**, but overlaid with a 3D representation of actin from the whole tomogram. **d**, Volume in **c** seen from the side. **e**, Slice through the average volume of sporozoite F-actin. **f**, Isosurface representation of the volume in **e** fitted with the molecular model of PDB 6TU4^15^. **g**, Size distribution of pellicular F-actin at different subcellular regions. Filaments that were fully contained within tomograms are shown in black, whereas those that were cut off by FIB milling are shown in grey. Bars represent medians. **h**, Cartoon representation of a sporozoite cell with colours corresponding to structures labelled in **a** and dotted lines showing lamella orientations for cells 1 and 2 in **b** and **c**.

We first analysed F-actin in the pellicular space with a view to understand the types of F-actin dynamics needed for rapid motility. At the apical pole, we observed several filaments in close proximity to the preconoidal rings (Fig. 2). Recent observations in related apicomplexans *Cryptosporidium parvum* and *Toxoplasma gondii* suggest that pellicular actin is nucleated at the preconoidal rings, implying that this is a consistent apicomplexan feature^11,12^. However, in *C. parvum* and *T. gondii*, the channeling of filaments into the pellicular space is dependent on extrusion of the conoid - a structural feature that is missing in *Plasmodium* sporozoites.

**Fig. 2:**
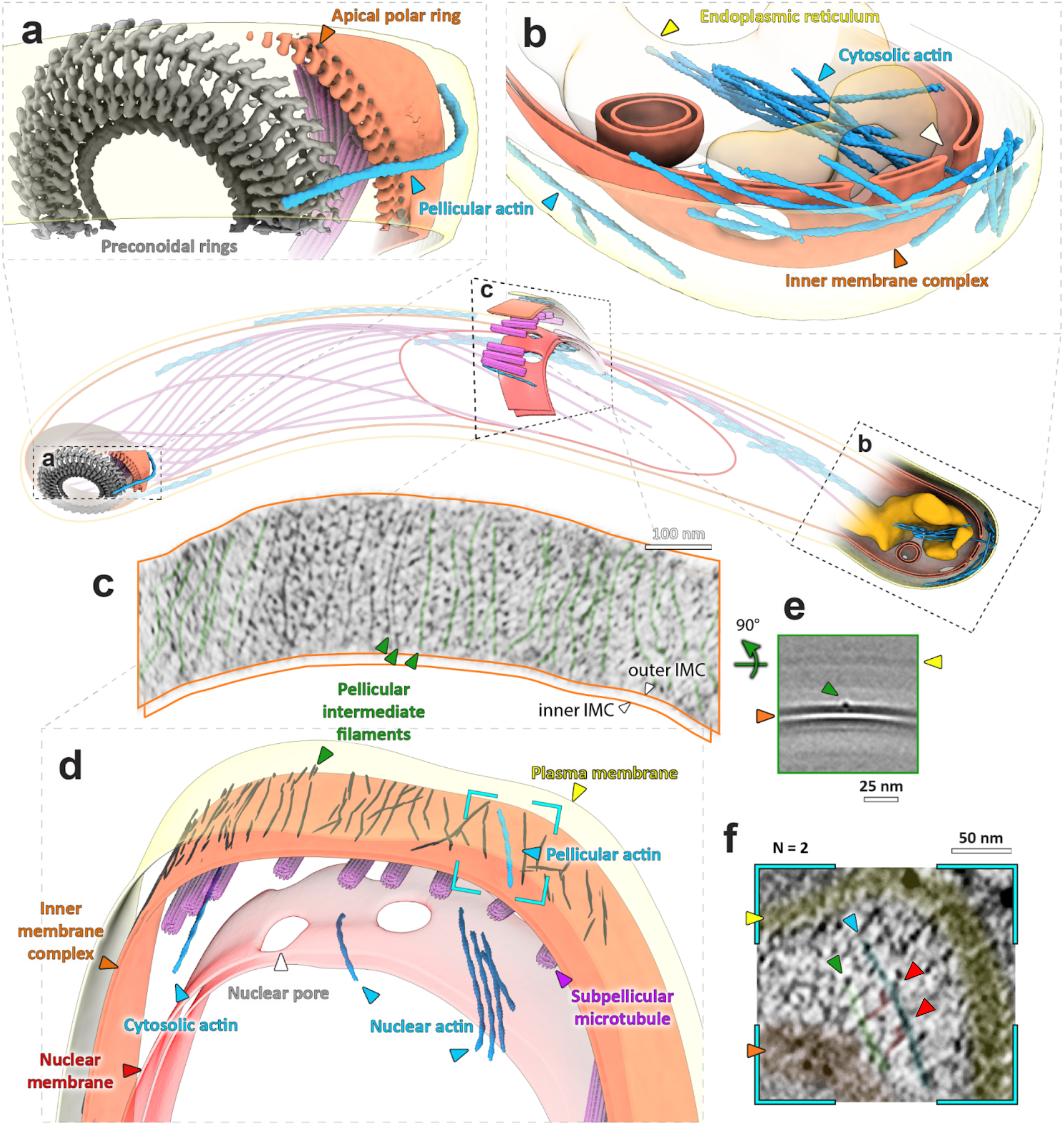
Visualising discrete actin populations within the 3D cellular context of motile sporozoites. Central cartoon represents the approximate position of volumes **a-c** within a cell. **a**, Apical end of showing a single actin filament being nucleated at the pre-conoidal rings (grey). **b**, Basal end of a sporozoite, showing cytoplasmic actin bundles as well as a build-up of pellicular actin. White arrowhead indicates an actin filament going through a basal pore. Shown is also an invagination of the IMC (observed in three cells). See slice through this tomogram in Fig. S3c. **c**, A lateral section showing nuclear, cytosolic and pellicular F-actin. Dark lines on the outer surface of the IMC represent pellicular intermediate filaments (PIFs). **d**, A slice along the surface of the outer leaflet of the outer IMC membrane. Some filaments have been highlighted in green. **e**, A slice through the average volume of PIFs showing their position relative to the IMC membranes. **f**, A slice through the tomogram (position shown in **c**) showing a pellicular actin filament (blue) connected to a PIF by two densities (red) consistent with myosin dimensions (observed in two different locations).

Analysis of the length distribution of pellicular F-actin shows two distinct length regulatory steps. (i) As filaments are transported down the cell towards the basal pole, they increase in length (Fig. 1g) and (ii) once at the basal pole, actin filaments are disassembled into shorter filaments (**F**ig. 1b-d,g). Short filaments at the basal pole supports a recently proposed model of severing mediated by coronin, an actin binding protein located at the basal end of gliding *P. berghei* sporozoites^13^, likely through recruitment of actin depolymerizing factors^14^. The shorter filaments accumulate at the basal pole, and in some cases form a one-filament deep shell, suggesting that actin disassembly may be rate-limiting (Fig. 1b,c). As actin disassembly occurs within the restricted pellicular space, a local actin monomer gradient likely forms within this space. The path of glideosome actin filaments towards the basal pole, may therefore be up an actin monomer gradient. The age of the filament together with this gradient could account for the elongation we observe as filaments move down the cell, prior to their disassembly at the basal pole. Localised pools of actin-stabilising proteins may also contribute to the length increase we observe.

The IMC in sporozoites has very few discontinuities, apart from at the basal pole which was dotted with ∼25 nm diameter pores (Fig. 2b, S3). We observed filamentous actin protruding through some basal pores, suggesting that filamentous actin can pass through into the cytoplasm (Fig. 2b, S3c, Video 1). It is therefore possible that these pores could facilitate more efficient exchange of G- and F-actin between the pellicular space and the rest of the cell. This would recycle actin and lower the concentration of actin at the rear of the sporozoite (up to 60 mM local F-actin concentration 100 nm from the basal pole end) and thus could allow for more efficient gliding. Strikingly, we observed few pores of the same size in other subcellular locations, indicating that their location at the basal pole is regulated, facilitating the directional flow of actin – into the pellicular space only via the preconoidal rings, and into the cytoplasm only via the basal pores. This is likely a conserved apicomplexan feature as similar pores were observed in *C. parvum*^11^, while *T. gondi* has a large opening in the IMC at the basal pole – both of which could facilitate directional actin flow. It would be reasonable to speculate that the posterior/basal polar ring complex^16^ could be involved in the localisation of the pores. Notably, we have not observed a ringlike structure at the basal pole, but rather a thick amorphous layer, which we refer to as basal polar matrix (Fig. S3c).

While examining the glideosome’s local environment, we noticed a network of thin filaments reinforcing the outer IMC membrane (membrane surface pointing towards the glideosome and plasma membrane) (Fig. 2c-f, 3b,c, Video 2). We refer to the filaments in *P. falciparum* as pellicular intermediate filaments (PIFs). PIFs were always oriented parallel to the long parasite axis and each other with a spacing of approximately 20 nm (∼5 - 40 nm) between them, with few clear cross-links. A SVA of PIFs highlighted that they are embedded in the outer IMC membrane (Fig. 3c). In two cellular locations we observed glideosomal actin filaments bound by densities consistent with myosin heads, with tails leading to PIFs in the IMC (Fig. 2f). The observed features prompted us to hypothesise that PIFs are structural elements that allow the force generated by the glideosome to be distributed along the entire IMC. PIFs bear no ultrastructural similarity to IMC surface filaments (IMCFs) observed in *C. parvum*, which form tracks to guide actin filaments down the cell^11^. Although it has been suggested that the gliding machinery is directly connected to subpellicular microtubules (SPMTs)^17^, with our current approach, we see no direct connection between PIFs, actin and SPMTs through the IMC membranes (Fig. 3). Although we observe homogenous ∼4 nm globular proteins that span the IMC membranes, and which we speculated may link the glideosome/PIFs to the SPMTS, these densities are not statistically associated with PIFs indicating no interaction (Fig. 3b,d).

**Figure 3.**
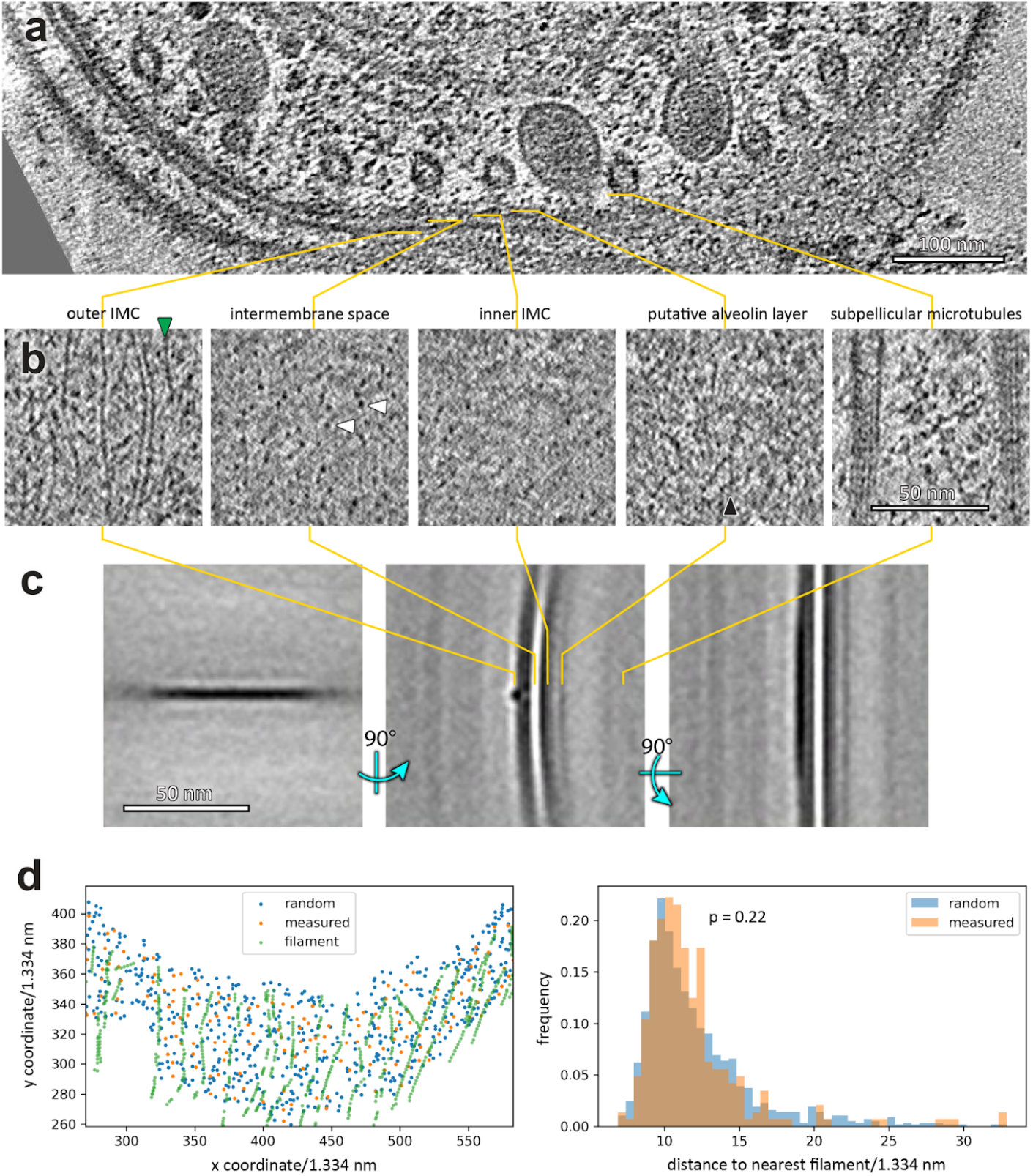
Analysis of the glideosome’s local environment; the pellicular space. a, A slice through a tomogram (also in Fig. 2c) highlighting the pellicle ultrastructure. **b**, Slices through tomogram shown in **a**, oriented parallel to the inner membrane complex (IMC) at indicated relative positions. We observed a large number of globular particles spanning the space between the two IMC membranes, with a homogeneous size distribution (∼ 4 nm diameter). **c**, Orthogonal sections through an average volume of ∼3000 PIF particles. **d**, We measured the distance distribution of the intermembrane particles to the nearest PIF to determine whether these may be directly interacting. However a randomly distributed set of coordinates (on a surface defined by the measured intermembrane particle coordinates, left) has the same distribution of distances (right), indicating that no interaction takes place.

Apart from motility, F-actin is implicated in multiple cellular roles including intracellular transport, transcriptional regulation and cell structural support. We therefore analysed the populations of F-actin that we observed in compartments other than the pellicular space.

We observed numerous examples of F-actin in the cytoplasm of sporozoites (Fig. 2b.d). This was found primarily as individual filaments, with some bundles (10 individual filaments and 4 bundles consisting of 2-8 filaments). F-actin was not associated in any obvious pattern with any subcellular element, but was typically oriented along the parasite’s apico-basal axis.

What was most surprising was the large amounts of F-actin we observed within the nucleus (Fig. 1a, 2d, Videos 2,3,4). In fact, the nucleus contains the majority of observed F-actin in sporozoites: approximately 60% (controlled for subcellular areas imaged in tomograms). It was found predominantly in bundles of ∼3-8 filaments with at least one filament less than 10 nm from the nuclear membrane (global median distance of all filaments was 21 ± 11 nm, Fig. 1a). Whether this is due to specific binding to a component of the membrane or due to marginalisation is not clear. While there have been many reports of the presence of nuclear actin in different organisms, this is to our knowledge, the first direct evidence of nuclear F-actin in the absence of staining or stabilising agents in any organism.

Indeed, actin chromobody signals have indicated F-actin accumulation near to the nucleus (in ∼20% of sporozoites) during motility and invasion of apicomplexan parasites, suggesting that a nuclear actin cage facilitates efficient invasion and/or protects the nucleus from damage when the parasite undergoes constriction^8,9^. However, here we have observed extensive bundles within the nucleus itself (Fig. 1a and 2d). Whether this simply serves as an intra-nuclear protective structure or fulfils additional molecular roles, such as mechanosensing, in the nucleus requires further investigation.

Actin is key for the transmission of the malaria parasite. By dissecting the path of F-actin within the glideosome we provide new mechanistic understanding of the dynamics of filament transport down the cell, severing at the basal pole and transfer and recycling via IMC pores. The optimal structural preservation and high resolution *in situ* data presented here will provide a framework for integrating future findings from other, more reductionist methods, in order to gain a complete understanding of the malaria parasite’s actin cytoskeleton in motility, in the cytoplasm and in the nucleus.

## Materials and Methods

### Obtaining sporozoites

*P. falciparum* sporozoites (strain: NF54-ΔPf47-5’csp-GFP-Luc: expressing a GFP-Luciferase fusion protein under the control of the csp promoter, genomic integration, no selection marker) were prepared at TropIQ (Nijmegen, Netherlands). Gametocytes were fed to 2 day old female *Anopheles stephensi* mosquitoes. Mosquito infection was confirmed 7 days post feeding by midgut dissection. At 7 days post infection, mosquitoes received an extra non-infectious blood meal to boost sporozoite production. Two weeks post infection, sporozoites were isolated using salivary gland dissection and shipped at room temperature in Leibovitz medium with 10% heat inactivated human serum.

### Cryo-grid preparation

*P. falciparum* sporozoites were checked under the fluorescent microscope and then diluted 1:4 into RPMI medium (without phenol red). 3 μl of parasites were applied onto a freshly plasma-cleaned UltrAufoil R1.2/1.3 300 mesh EM grid (Quantifoil) in a humidity controlled facility. Excess liquid was manually back-blotted and grids were plunged into a reservoir of ethane/propane using a manual plunger. Grids were stored under liquid nitrogen until imaging.

### Cryo FIB milling

Grids were clipped into autogrids modified for FIB preparation^17^ and loaded into either an Aquilos or an upgraded Aquilos2 cryo-FIB/SEM dual-beam microscope (Thermofisher Scientific). Overview tile sets were recorded using MAPS software (Thermofisher Scientific) before being sputter coated with a thin layer of platinum. Good sites with parasites were identified for lamella preparation before the coincident point between the electron beam and the ion beam was determined for each point by stage tilt. Prior to milling, an organometallic platinum layer was deposited onto the grids using a GIS (gas-injection-system). Lamellae were milled manually until under 300 nm in a stepwise series of decreasing currents. Milling was performed at the lowest possible angles to increase lamella length in thin cells. Finally, polishing of all lamella was done at the end of the session as quickly as possible but always within 1.5 h to limit ice contamination from water deposition on the surface of the lamellae. Before removing the samples, the grids were sputter coated with a final thin layer of platinum. Grids were stored in liquid nitrogen for a maximum of 2 weeks before imaging in the TEM.

### Tilt-series collection

Cryo-EM FIB-milled grids were rotated by 90° and loaded into a Titan Krios microscope (Thermofisher) equipped with a K3 direct electron detector and (Bio-) Quantum energy filter (Gatan). Tomographic data was collected with SerialEM with the energy-selecting slit set to 20 eV. Datasets were collected using the dose-symmetric acquisition scheme at a ± 65° tilt range with 3° tilt increments. For all datasets, 5-10 frames were collected and aligned on the fly using SerialEM and the total fluence was kept to less than 120 e−Å^2^. Defoci between 3 and 8 μm underfocus were used to record the tilt series’.

### Tomogram reconstruction

Frames were aligned on the fly in SerialEM^18^; CTF estimation, phase flipping and dose-weighting was performed in IMOD^19^. Tilt-series’ were aligned in IMOD either using patch-tracking or by using nanoparticles (likely gold or platinum) on lamella surfaces as fiducial markers. Tomograms were binned 4x and filtered in IMOD or by using Bsoft^20^.

### Subvolume averaging

Subvolume averaging was performed using PEET^21^ as described previously^22^. Model processing was done using TEMPy^23^, Scipy^24^, Scikitlearn^25^, Matplotlib26 and Numpy^27^ in Python 3. Initial models were generated manually by picking line segments using pairs of IMOD model points and then interpolating particles at 1 voxel (1.3 nm) increments. The initial Y axes were aligned with the line segments and initially Y axis rotation angles were randomised. The initial reference was generated by averaging particles with the starting orientations, thus generating a featureless cylinder. A small subset of particles (∼700) were refined to create a reference with F-actin features which was then used for alignment of ∼ 70k initial positions. Duplicate and low scoring particles were removed. In order to improve model completeness and allow separation of particles into two independent halves, the subvolume positions were then fitted to a spline-smoothed helical model allowing for small variation in helical pitch (Figure S2). Subvolume positions were then generated based on the best fitting model parameters. These were split into two halves and aligned independently. Overlapping particles between the two halfmaps were removed before generating final half-maps. Fourier Shell Correlation was measured using Bsoft, suggesting 27 Å resolution at the 0.143 cutoff. Particles from the two half-datasets (11487 total) were then combined and aligned together. The final volume was sharpened using Bsoft with an arbitrarily chosen B-factor of -3000 for fitting and visualisation.

### Segmentation and visualisation

Membrane segmentation was performed in IMOD, using drawing tools followed by linear interpolation. These were then resampled using open3d to achieve an isotropic coordinate distribution, which were then used to generate a volume using IMOD imodmop. F-actin, microtubules, apical polar ring and preconoidal rings were backplotted: average volumes were placed into 3D volumes using coordinates determined by SVA. Actin and microtubule models were smoothed for backplotting. Surface visualisation was performed using UCSF ChimeraX or open3d. Volume sections were visualised using IMOD 3dmod. Plots were generated using Matplotlib.

### Length measurements

Filament lengths for comparison of nuclear, cytosolic and pellicular filament lengths were derived from helical models based on subvolume averaging positions (see above). Filament lengths for comparison of apical, lateral and basal pellicular filament lengths were measured manually using 3dmod.

### F-actin concentration

The number of actin subunits in observed F-actin was estimated from subvolume averaging (15,058) and manual length measurements (17165, assuming 38 nm per 13 subunits). The subvolume averaging-derived value is likely an underestimate due to cross-correlation based particle cleaning; it is the number of particles after the first alignment step of the two independent datasets. The two estimates were used to calculate the experimental error, expressed as standard deviation. The total observed volume of 29 tomograms with an average thickness of 244 nm was 4.2 × 10-17 m^3^, of which cells made up approximately 7/12. 9.7 × 10^−19^ mol in 2.4 × 10^−14^ L corresponds to 4.0 × 10^−5^molL^-1^

## Supporting information

Vieo 4

## Acknowledgements

We thank Lindsay Baker for helpful discussions and Carolyn Moores for her continued support and critical reading of the manuscript. Thank you to the CSSB EM facility team for their support. We gratefully acknowledge funding by HFSP long-term postdoctoral fellowship LT000024/2020-L (JLF), Infrastructures for the control of vector-borne diseases (Infravec2) funded by the EU’s Horizon 2020 programme (grant agreement No 731060) (JLF), Wellcome Career Development award 227774/Z/23/Z (JLF), LOEWE Centre DRUID (“Novel Drugs Targets against Poverty-related and Neglected Tropical Infectious Diseases”) within the Hessian Excellence Program (RGD). For the purpose of open access, the author has applied a Creative Commons Attribution (CC BY) licence to any Author Accepted Manuscript version arising. Biorxiv template from:www.github.com/chrelli/bioRxiv-word-template

## Author contributions

VP and JLF designed the study and experiments. JLF generated samples and acquired data. JLF and VP processed and analysed data. VP, JLF, RGD, DV, KG performed critical analysis of findings. JLF, VP, RGD wrote the manuscript. VP generated figures.

## Data availability

All data are available on request. Subvolume average was deposited on the EMDB.

## Code availability

Scripts are available on request.

## Competing interests

The authors declare no competing interests.

## Notes

### Competing Interest Statement

The authors have declared no competing interest.

### Summary of Updates

This manuscript has been revised to add new discussion points, additional references and one figure has moved from supplemental to the main text. We have also added four movies which help with understanding the data.

## References

1. Cowman, A. F., Healer, J., Marapana, D. & Marsh, K. Malaria: Biology and Disease. Cell 167, 610–624 (2016).

2. Amino, R. et al. Quantitative imaging of Plasmodium transmission from mosquito to mammal. Nat. Med. 12, 220–224 (2006).

3. Heintzelman, M. B. Gliding motility in apicomplexan parasites. Semin. Cell Dev. Biol. 46, 135–142 (2015).

4. Schmitz, S. et al. Malaria Parasite Actin Filaments are Very Short. J. Mol. Biol. 349, 113–125 (2005).

5. Vahokoski, J. et al. Structural Differences Explain Diverse Functions of Plasmodium Actins. PLoS Pathog. 10, e1004091 (2014).

6. Lu, H., Fagnant, P. M. & Trybus, K. M. Unusual dynamics of the divergent malaria parasite PfAct1 actin filament. Proc. Natl. Acad. Sci. U. S. A. 116, 20418–20427 (2019).

7. Kumpula, E.-P. et al. Apicomplexan actin polymerization depends on nucleation. Sci. Rep. 7, 12137 (2017).

8. Yee, M., Walther, T., Frischknecht, F. & Douglas, R. G. Divergent Plasmodium actin residues are essential for filament localization, mosquito salivary gland invasion and malaria transmission. PLOS Pathog. 18, e1010779 (2022).

9. Del Rosario, M. et al. Apicomplexan F-actin is required for efficient nuclear entry during host cell invasion. EMBO Rep. 20, e48896 (2019).

10. Tosetti, N., Dos Santos Pacheco, N., Soldati-Favre, D. & Jacot, D. Three F-actin assembly centers regulate organelle inheritance, cell-cell communication and motility in Toxoplasma gondii. eLife 8, e42669 (2019).

11. Martinez, M. et al. Origin and arrangement of actin filaments for gliding motility in apicomplexan parasites revealed by cryo-electron tomography. Nat. Commun. 14, 4800 (2023).

12. Dos Santos Pacheco, N. et al. Conoid extrusion regulates glideosome assembly to control motility and invasion in Apicomplexa. Nat. Microbiol. 7, 1777–1790 (2022).

13. Bane, K. S. et al. The Actin Filament-Binding Protein Coronin Regulates Motility in Plasmodium Sporozoites. PLOS Pathog. 12, e1005710 (2016).

14. Douglas, R. G. et al. Inter-subunit interactions drive divergent dynamics in mammalian and Plasmodium actin filaments. PLOS Biol. 16, e2005345 (2018).

15. Vahokoski, J. et al. High-resolution structures of malaria parasite actomyosin and actin filaments. PLOS Pathog. 18, e1010408 (2022).

16. De Niz, M. et al. Progress in imaging methods: insights gained into Plasmodium biology. Nat. Rev. Microbiol. 15, 37–54 (2017).

17. Harding, C. R. et al. Alveolar proteins stabilize cortical microtubules in Toxoplasma gondii. Nat. Commun. 10, 401 (2019).

18. Schaffer, M. et al. Cryo-focused Ion Beam Sample Preparation for Imaging Vitreous Cells by Cryo-electron Tomography. Bio-Protoc. 5, e1575 (2015).

19. Mastronarde, D. N. Dual-Axis Tomography: An Approach with Alignment Methods That Preserve Resolution. J. Struct. Biol. 120, 343–352 (1997).

20. Kremer, J. R., Mastronarde, D. N. & McIntosh, J. R. Computer visualization of three-dimensional image data using IMOD. J. Struct. Biol. 116, 71–76 (1996).

21. Heymann, J. B. Bsoft: Image and Molecular Processing in Electron Microscopy. J. Struct. Biol. 133, 156–169 (2001).

22. Heumann, J. M., Hoenger, A. & Mastronarde, D. N. Clustering and variance maps for cryo-electron tomography using wedge-masked differences. J. Struct. Biol. 175, 288–299 (2011).

23. Ferreira, J. L. et al. Variable microtubule architecture in the malaria parasite. Nat. Commun. 14, 1216 (2023).

24. Cragnolini, T. et al. TEMPy2: a Python library with improved 3D electron microscopy density-fitting and validation workflows. Acta Crystallogr. Sect. Struct. Biol. 77, 41–47 (2021).

25. Virtanen, P. et al. SciPy 1.0: fundamental algorithms for scientific computing in Python. Nat. Methods 17, 261–272 (2020).

26. Buitinck, L. et al. API design for machine learning software: experiences from the scikit-learn project. Preprint at 10.48550/arXiv.1309.0238 (2013).

27. Hunter, J. D. Matplotlib: A 2D Graphics Environment. Comput. Sci. Eng. 9, 90–95 (2007).

28. Harris, C. R. et al. Array programming with NumPy. Nature 585, 357–362 (2020).

