## Supplemental figures for "Molecular architecture of glideosome and nuclear F-actin in *Plasmodium falciparum*"

### Supplementary Material:

**Video 1:** Movie moving through a tomogram showing two *Plasmodium falciparum* sporozoite basal poles. Basal pores are seen with an example of an actin filament sitting within one pore.

**Video 2:** Movie of a tomogram that has been rotated to move through all regions of the pellicle. Includes the plasma membrane, PIFs, pellicular actin, IMC membranes, subpellicular microtubules, nuclear F-actin and a nuclear pore complex.

**Video 3:** Movie moving through a tomogram showing the nucleus of a *Plasmodium falciparum* sporozoite with a bundle of nuclear actin.

**Video 4:** Movie moving through a tomogram showing the nucleus of a *Plasmodium falciparum* sporozoite, with nuclear pores and nuclear filamentous actin. The apical pole of a neighbouring sporozoite is also seen.

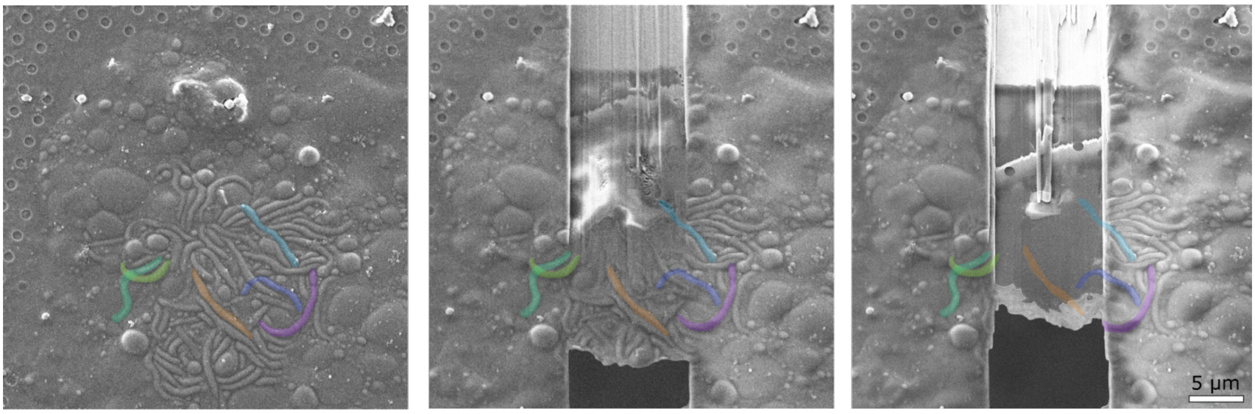


**Supplementary Figure 1. FIB-milling sporozoites. Left:** A pile of sporozoites in the SEM. A few individual sporozoites are coloured consistently in all three images. **Middle:** SEM image during the FIB-milling process. **Right:** SEM image of the final polished lamella.

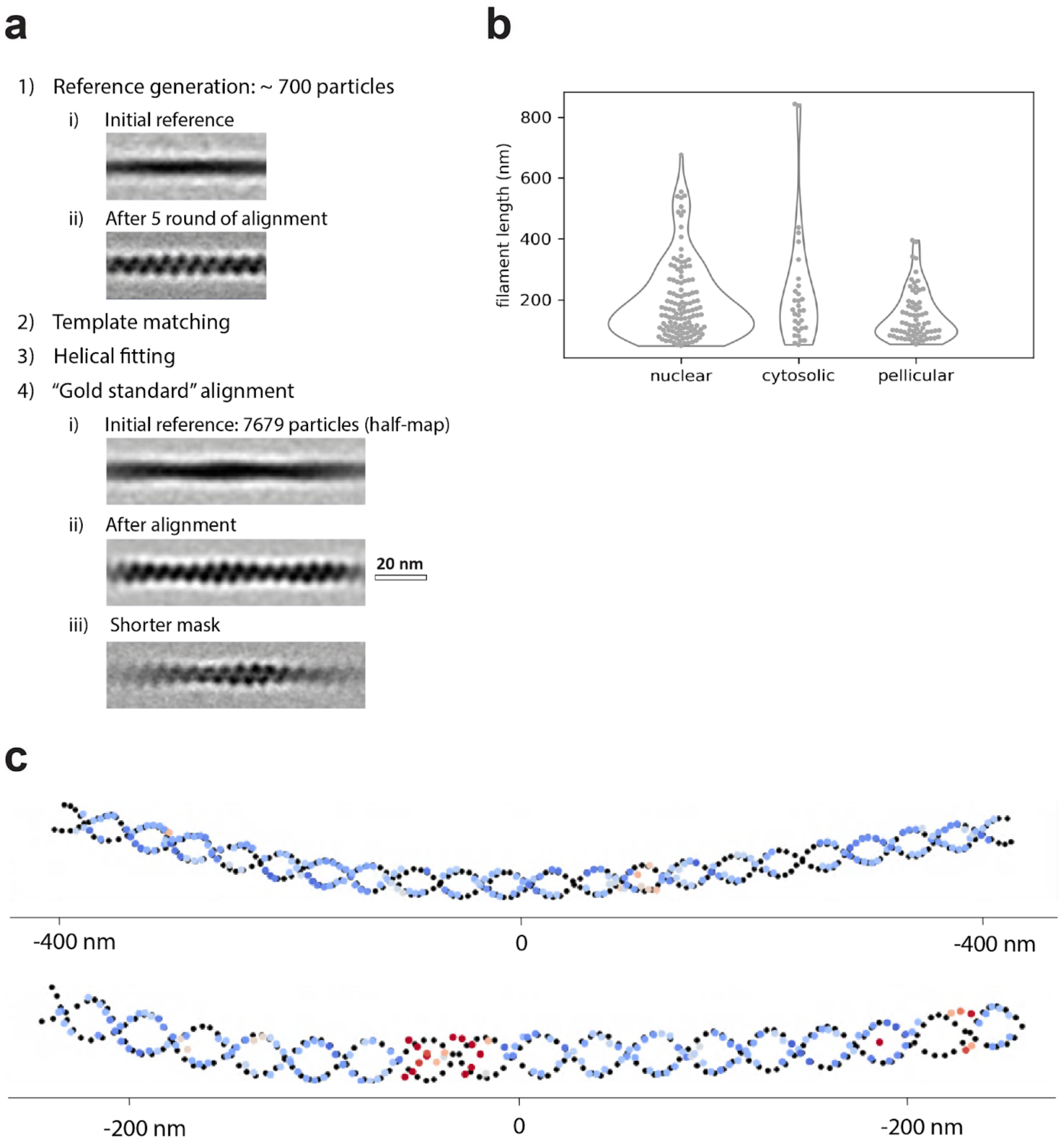


**Supplementary Figure 2. Subvolume averaging of actin. a,** A rough outline of the workflow used for subvolume averaging of actin, with some reference volumes shown. **b,** Size distribution of F-actin in indicated subcellular locations derived from subvolume averaging coordinates. Note that especially longer filaments were frequently truncated during FIB milling causing the distribution to be skewed towards shorter sizes. Two-sided t-test p-values are 0.4 and $1 \times 10^{-4}$ comparing nuclear and cytosolic, and nuclear and pellicular, respectively. Caution should be used interpreting the significance of the difference due to some filaments being truncated. **c,** Diagnostic plots of helical fitting of two longer filaments. Black dots show best fit positions of actin subunits. Overlaid are positions measured by subvolume averaging coloured from blue to red based on distance to the nearest model point.

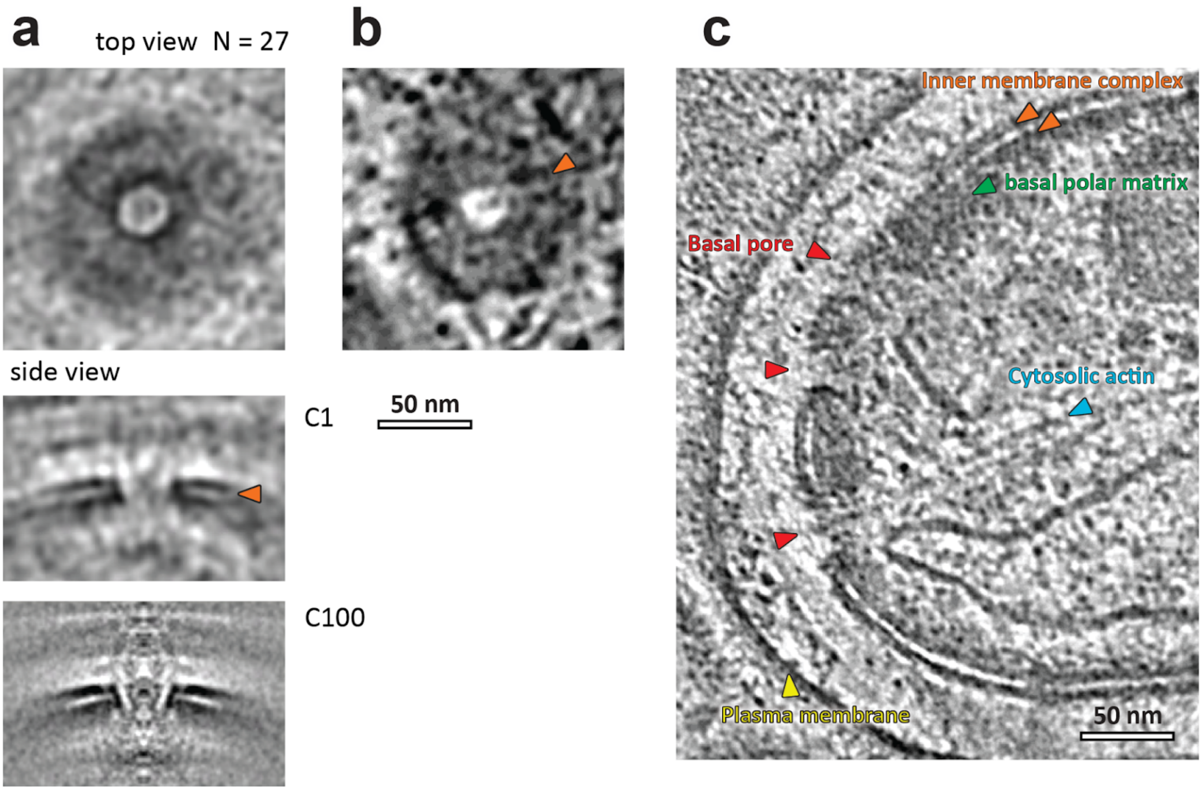


**Supplementary Figure 3: Basal pores form 25 nm connections between the cytoplasm and pellicular space. a,** Subvolume average of 27 particles. No symmetry is evident at this resolution. **b,** A tangential slice through a single pore. **c,** Slice through a basal end of a sporozoite (also in Fig. 2b) showing three basal pores.
